# Iron and Lipocalin-2 Modulate Cellular Responses in the Tumor Micro-environment of Pancreatic Ductal Adenocarcinoma

**DOI:** 10.1101/2020.01.14.907188

**Authors:** Valentina Pita-Grisanti, Andrew W. Dangel, Kristyn Gumpper, Andrea Ludwig, Olivia Ueltschi, Xiaokui Mo, Maciej Pietrzak, Amy Webb, Rosa F. Hwang, Madelyn Traczek, Niharika Badi, Zobeida Cruz-Monserrate

## Abstract

Pancreatic ductal adenocarcinoma (PDAC) is a highly metastatic disease with poor outcomes. Iron is known to signal cellular responses, and its levels are regulated by lipocalin-2 (LCN2) expression, a PDAC pro-tumorigenic molecule. However, how iron and LCN2 function in PDAC is unclear. Here we demonstrate that iron levels regulate PDAC cell proliferation, invasion, expression of epithelial to mesenchymal tumor markers, and pro-inflammatory cytokines. Iron chelation increased the expression of the LCN2 receptor *SLC22A17* in pancreatic stellate cells and the anti-metastatic gene *NDRG1* in PDAC cells. Deletion of *Lcn2* in mouse tumor cells modulated the expression of genes involved in extracellular matrix deposition and cell migration. Moreover, cellular iron responses were dependent on the *Kras* mutation status of cells, and *LCN2* expression levels. Deletion of *Lcn2* expression in PDAC suggests a protective role against metastasis. Thus, iron modulation and LCN2 blockade could serve as potential therapeutic approaches against PDAC.

## Introduction

Pancreatic ductal adenocarcinoma (PDAC) is currently the third leading cause of cancer-related deaths in the United States, mostly due to the lack of early detection methods, late diagnosis, and limited treatment options (1). Poor prognosis results from late stage diagnosis, low eligibility for tumor resection, increased resistance to current therapies, high cancer cell proliferation rates, and elevated incidence of metastases (2–4).

PDAC progression has been associated with increased expression of pro-inflammatory molecules such as lipocalin-2 (LCN2) (5). LCN2 is a secreted glycoprotein involved in the innate immune response that is upregulated in many cancers including PDAC (5, 6). Our group has previously shown that LCN2 expression is highly elevated in the blood of PDAC patients and in PDAC tumor cells, inducing inflammation by modulating the secretion of pro-inflammatory cytokines in human pancreatic stellate cells (HPSCs) in the PDAC tumor microenvironment (TME) (5). Moreover, depletion of LCN2 in mice extends survival and delays PDAC tumor growth (5). LCN2 chelates divalent and trivalent iron via siderophore binding that controls cellular iron uptake and apoptosis (7–9), for this reason, iron modulation could be essential in regulating the mechanisms by which LCN2 contributes to the tumorigenesis of PDAC.

Cellular iron levels control metabolism, proliferation, DNA synthesis, and cell death of both normal and neoplastic cells (10–14). Cancer cells increase iron uptake and expression of the transferrin receptor, known to transport iron into the cells (15). Moreover, iron metabolism-related pathways are used as prognostic indicators of various cancers (16–18). Increased iron levels are involved in the epithelial-mesenchymal transition (EMT) of cancer cells, leading to increased metastasis (10, 11, 18–26). Since PDAC cells undergo EMT, decreasing iron levels could potentially inhibit EMT in PDAC (18, 23, 27–29). Iron chelators, such as deferoxamine (DFO), are used to reduce iron levels and have been effective in the treatment of iron overload diseases like hemochromatosis (30). In leukemia, breast, and colorectal cancers, DFO treatment inhibits cell growth and promotes apoptosis (31–33). Iron also regulates the expression of genes involved in metastasis. Among these, high levels of iron downregulate the expression of the N-myc downstream-regulated gene1 (NDRG1). NDRG1 is known to suppress metastasis and inhibit EMT in several cancers, including PDAC (34), and it is associated with the differentiation state of PDAC cells, with well-differentiated cells expressing higher levels of NDRG1 (35).

Given the increasing evidence suggesting a link between LCN2, iron, and PDAC tumorigenesis, here we investigated the effects of iron level modulation and LCN2 expression on cell proliferation, invasion, and expression of various inflammatory-related cytokines on PDAC cancer and stromal-derived cells. Moreover, we assessed whether LCN2 depletion from cancer cells resulted in gene expression changes related to EMT and metastasis.

## Results

### Iron levels regulate proliferation of PDAC and pancreatic stellate cells

Iron is known to modulate cell proliferation (12, 13). Therefore, we tested whether modulating iron levels *in vitro* by adding or chelating iron would affect the proliferation of PDAC and pancreatic stellate cells. Since iron responses are known to be dependent on the *Kras* mutation status of cancer cells (36) we selected to study PANC1 (mutant for Kras) and BXPC3 (wild-type for Kras) human PDAC cells (37). We treated human PDAC cells, mouse PDAC cells (KPC) and human PSC (HPSC) cells with increasing concentrations of iron (ferrous ammonium citrate, FAC). FAC treatments of 0.313 mM and 20 mM reduced proliferation of BXPC3, however FAC concentrations between 0.625 mM and 10 mM at 48 and 72 hours increased proliferation (**Figure 1**). In contrast, increased FAC treatments decreased the proliferation of PANC-1, KPC, and HPSC cells in a dose-dependent manner for which HPSCs were more sensitive to FAC (**Figure 1 and Figure 1 – figure supplement 1A**). To address whether iron chelation alone inversely regulated cell proliferation, cells were treated with the iron chelator DFO. At lower doses of DFO, we observed an increased trend towards proliferation for all cells with maximum proliferation at 48 hours, which was statistically significant only in HPSC **(Figure 1B, D, F, and Figure 1-figure supplement 1B**). In contrast, higher doses of DFO (12.5 µM, 25 µM, and 50 µM) resulted in decreased proliferation for all cells after 72 hours except for KPC cells **(Figure 1B, D, F, and Figure 1 – figure supplement 1B**). Therefore, cell proliferation is dependent on iron level modulation and suggests that iron levels could impact cells proliferation in a Kras-dependent matter in PDAC.

**Figure 1.**
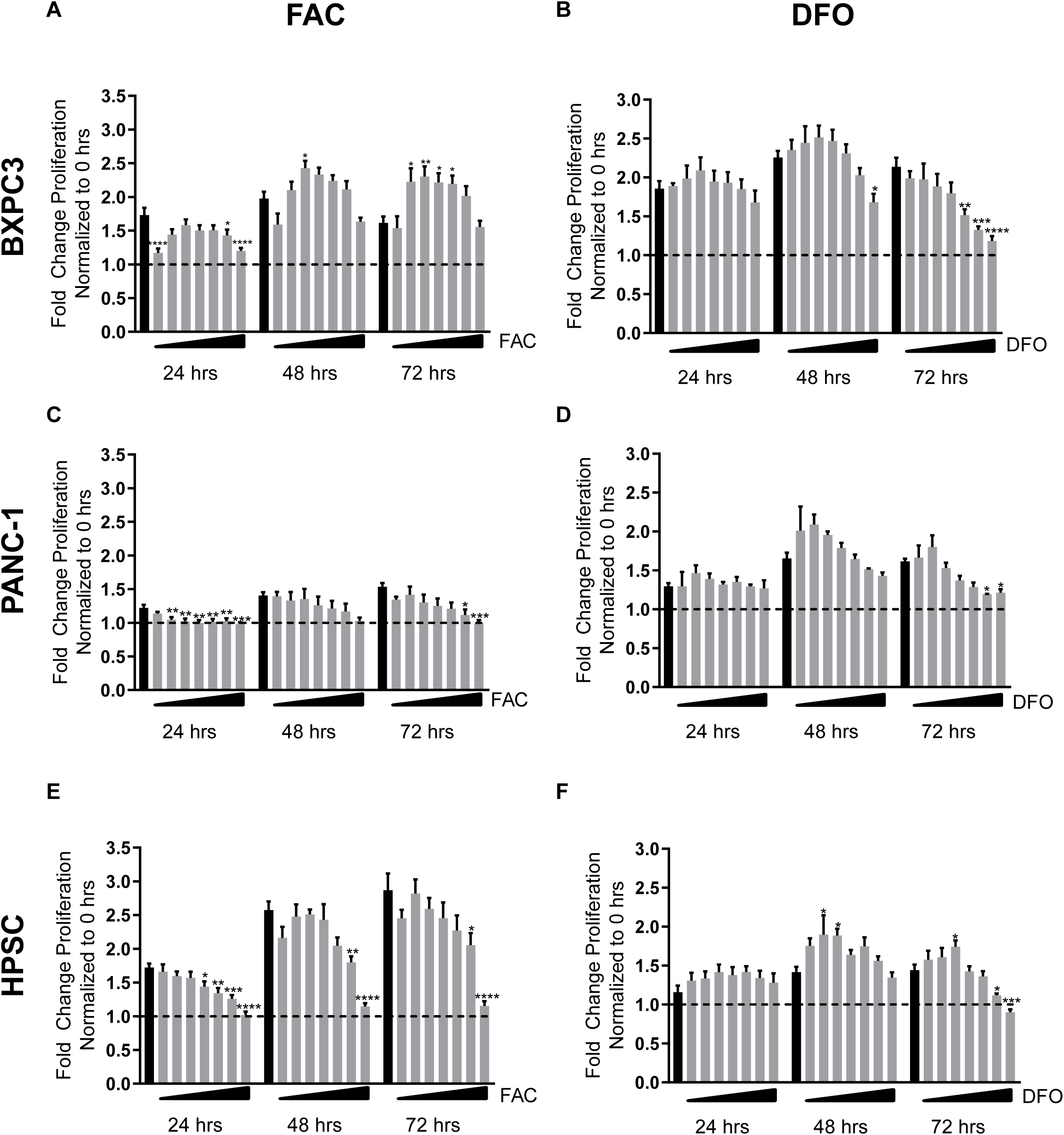
Iron and DFO treatments regulate cell proliferation in the TME. (**A**), BXPC3, (**C**), PANC-1 and (**E**). HPSC were treated with, 0.313 mM, 0.625 mM, 1.25 mM, 2.5 mM, 5 mM, 10 mM, and 20 mM, FAC. (**B**) BXPC3, (**D**) PANC-1 and (**F**) HPSC were treated with 0.781 µM, 1.563 µM, 3.125 µM, 6.25 µM, 12.5 µM, 25 µM, and 50 µM DFO. Cell proliferation was measured using MTS following exposure to FAC or DFO over a range of 72 hours. Results were normalized to 0 hours, represented by a horizontal dashed line. Significance was assessed by one-way ANOVA comparison test. Bars represent mean ± SEM. *p≤0.05, **p≤0.01, ***p≤0.001 ****p≤0.0001. Sample size ranged from 3 to 12 replicates, each group. Black bars represent non-treated cells.

### Iron levels modulate the expression of pro-inflammatory and iron-transport genes in PDAC and pancreatic stellate cells

TME-associated inflammation can mediate tumor growth (38, 39). Therefore, we tested whether modulating iron levels could influence the expression of pro-inflammatory genes (*IL6* and *IL1β*), known to be regulated by the expression of LCN2, a PDAC-associated cytokine involved in cellular iron uptake (5). We also measured the expression of two iron-transport genes, ferritin heavy chain 1 (*FTH1*), and solute carrier family 22 member 17 (*SLC22A17),* (LCN2 receptor) to verify cellular iron storage, and iron-bound LCN2 transport into the cells respectively. BXPC3, PANC-1, and HPSC were treated with FAC at 150µM (a physiologically relevant dose of iron) (40) for 24 hours and showed elevated expression of *IL6*, *IL1β*, and *FTH1*, in the presence of FAC relative to control (**Figure 2)**. However, *SLC22A17* expression was not responsive to iron treatment in any of the cell lines tested **(Figure 2)**. In addition, we treated cells with 20 µM of DFO, and showed that expression of *IL6* and *IL1β* decreased in HPSC. Expression of *IL6* was not stimulated in PANC-1 and BXPC3 while *IL1β* increased in BXPC3 after iron chelation (**Figure 2B, D**). *FTH1* expression was reduced in PANC-1, while it was increased in HPSCs after DFO treatment **(Figure 2F)**. Moreover, S*LC22A17* expression was upregulated only in HPSCs after iron chelation **(Figure 2H),** which could be the result of an adaptation mechanism in response to low levels of iron in HPSCs, to preserve iron transport in the TME. Thus, increased iron levels seem to promote inflammation and iron transport in PDAC and HPSCs, but do not affect the expression of the LCN2 receptor. Iron chelation for the most part blocked some of those effects and specifically increased iron transport molecules in HPSCs. These results indicate that stromal cells of the TME respond differently to reduced iron levels than cancer cells.

**Figure 2.**
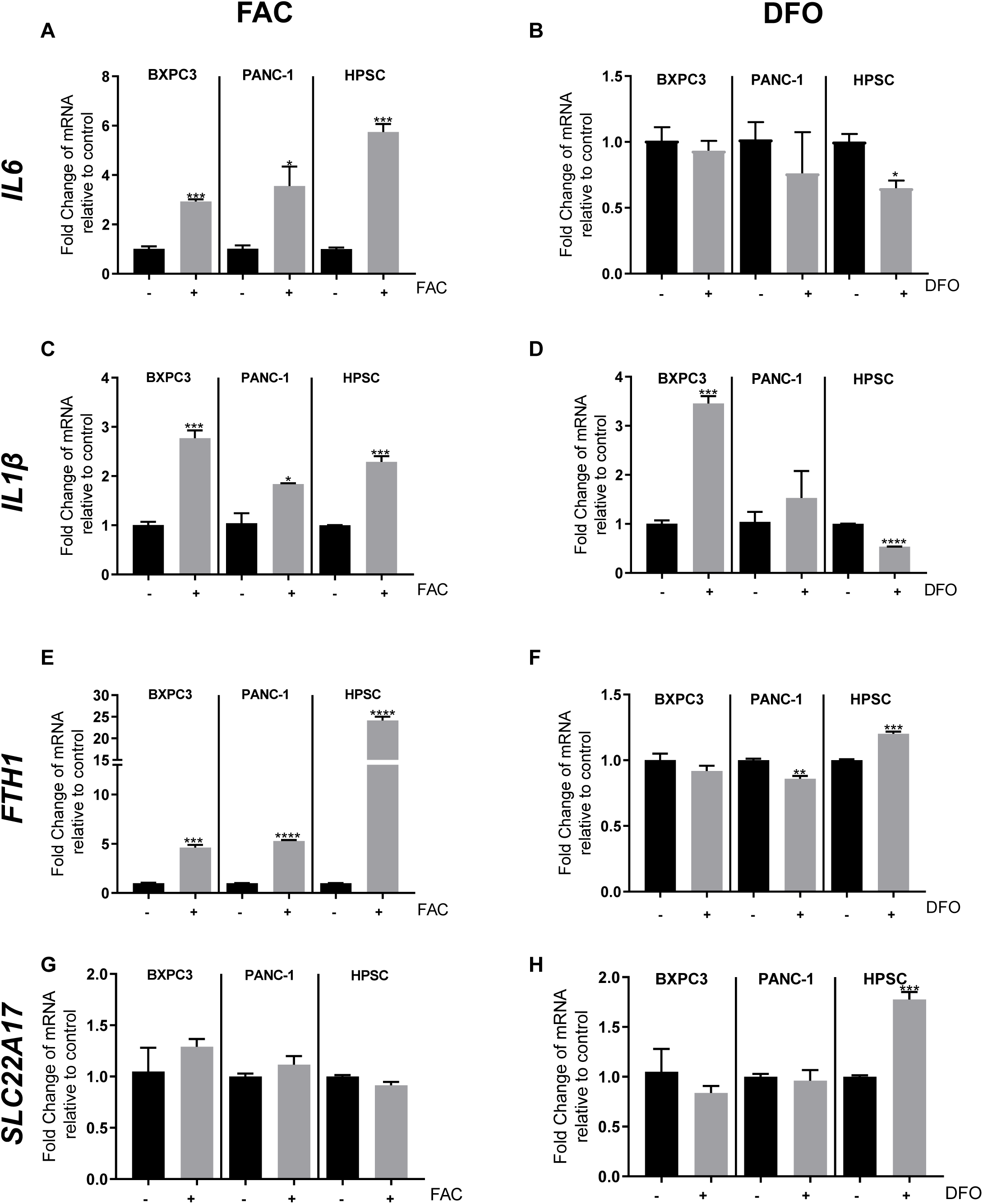
Iron and DFO levels modulate the expression of pro-inflammatory cytokines and iron-transport genes in the TME. Gene expression levels for (**A**) *IL6*; (**C**) *IL1β*; (**E**) *FTH1* and *SLC22A17* in BXPC3, PANC-1, and HPSC with and without iron treatment. Results are relative to 0 µM FAC (-), (+) denotes 150 µM FAC, maintained for 24 hours. Gene expression levels for (**B**) *IL6;* (**D**) *IL1β*; (**F**) *FTH1* and (**H**) *SLC22A17* in BXPC3, PANC-1 and HPSC with and without DFO treatment relative to 0 µM DFO (-), (+) denotes 20 µM DFO, maintained for 24 hours. Unpaired t-test was used to compare the groups. Bars represent mean ± SEM. *p≤0.05, **p≤0.01, ***p≤0.001 ****p≤0.0001. n=3 replicates.

### Iron treatment promotes EMT and cancer cell invasion of human PDAC cell lines in a Kras-dependent matter

Iron promotes changes in EMT which is known to precede invasion (10). Therefore, to understand whether iron levels modulate the EMT phenotype of PDAC cells, we examined cell morphology and expression of EMT markers as a result of iron treatment. Increased iron induced a mesenchymal morphology in BXPC3 that was not observed in PANC-1 cells **(Figure 3A)**. To confirm the morphological changes observed, classical EMT markers, *ZEB1*, *SNAI1*, and *TWIST* transcription factors, and the epithelial marker E-cadherin (*CDH1*) were measured after 48 hours of 20mM FAC treatment **(Figure 3B, C)**. In BXPC3 cells, expression levels of all the EMT gene markers were elevated after FAC treatments, while *CDH1* (an epithelial marker) expression was decreased, as expected for cells undergoing EMT. *TWIST* had the largest increased in gene expression after FAC treatment (5.5-fold increase) compared to control in BXPC3 cells. In PANC1 cells FAC did not induced expression of EMT markers, but it resulted in a 2.7-fold decrease in expression of *CDH1*. Moreover, iron chelation decreased the expression of EMT markers (*ZEB1, SNAI1 and TWIST*) in both BXPC3 and PANC-1 (**Figure 3 - figure supplement 1A, B**). Interestingly, DFO also decreased *CDH1* expression in PANC-1. These data suggest that iron chelation could inhibit iron-dependent EMT of cancer cells and might be dependent on the Kras mutations status of cells. To further assess how the iron-dependent EMT modulation regulates the invasive potential of BXPC3 and PANC-1, we measured invasion via transwell assay with a BME coated membrane. We showed that iron treatments significantly increased the invasion of BXPC3 cells at both 150µM and 20mM FAC. However, invasion was significantly increased only in PANC-1 cells after 150µM FAC and not 20mM FAC **(Figure 3D, E)**. These correlates with the increased expression of EMT markers observed in BXPC3 cells at 20mM FAC.

**Figure 3.**
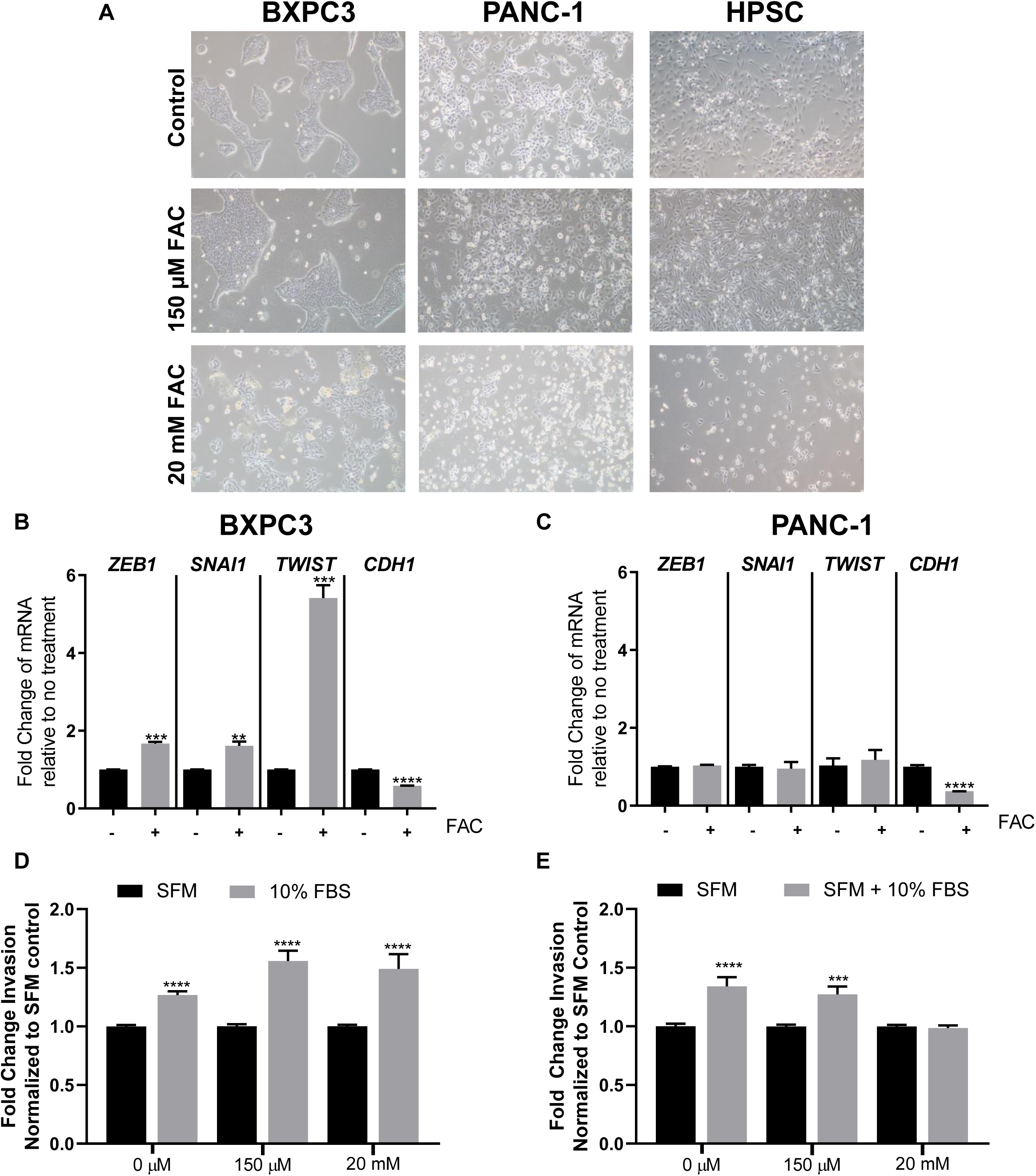
Increased iron levels upregulate genes involved in EMT and promotes an invasive phenotype in BXPC3. (**A**) Phase contrast images of BXPC3, PANC-1 and HPSC treated for 48 hours with 150 µM FAC or 20 mM FAC, compared to non-treated control. (**B**) Gene expression levels for *ZEB1*, *SNAI1*, *TWIST*, and *CDH1* in BXPC3 (**B**) and PANC-1 (**C**) relative to 0 mM FAC (-) treatment and 20 mM FAC (+). Treatments were maintained for 48 hours. Invasion assays for BXPC3 (**D**) and PANC-1 (**E**) with 150 µM FAC or 20 mM FAC treatments. Fold change relative to media no FBS for each treatment. Significance was assessed by unpaired t-test. Bars represent mean ± SEM, results are normalized to non-treated cells in SFM. *p≤0.05, **p≤0.01, ***p≤0.001 ****p≤0.0001. n=3 replicates for **B** and **C**, n=3-5 independent experiments for **D** and **E**.

### Iron regulates *NDRG1* expression which is inversely correlated with *LCN2* expression in PDAC

Iron is known to downregulate the expression of the iron related metastasis suppressor, N-myc downstream regulated gene 1 (NDRG1), and this downregulation is associated with increased proliferation and invasion of PDAC (34, 41, 42). Therefore, we measured the expression of *NDRG1* after treating BXPC3 and PANC-1 cells with 150μM and 20mM FAC. We showed that NDRG1 expression decreases 33-fold in BXPC3 and 22-fold in PANC-1 cells after FAC treatment **(Figure 4A)**. NDRG1 expression was considerably higher in BXPC3 compared to PANC-1 cells not treated with FAC. Moreover, iron chelation induced the expression of *NDRG1* in both BXPC3 and PANC-1 cells (**Figure 4B**). Our data suggests that iron chelation could decrease iron-induced metastasis via EMT and NDRG1 regulation. *NDRG1* expression is involved in cell line differentiation, with well differentiated PDAC cells expressing higher levels of *NDRG1* and poorly differentiated PDAC cells expressing lower levels or no *NDRG1* mRNA (35). Moreover, *NDRG1* expression was found to be negatively regulated by LCN2 in cholangiocarcinoma cells (43). Therefore, we assessed the expression of *LCN2* and *NDRG1* in multiple human PDAC cell lines and a HPSC cell line compared to a normal human pancreatic ductal epithelial cell line (HPDE). We showed that in general there was an inverse relationship between *NDRG1* expression and *LCN2* expression, except for PANC-1 and MIAPACA2, where both *NDRG1* and *LCN2* expression were low (**Figure 4C and 4E)**. Since these genes are involved in iron regulation, we also assessed the expression of the iron transport gene *FTH1* and showed that all cell lines had significantly lower expression levels than HPDE. **(Figure 3 - figure supplement 1C)**. To validate our findings in another model, we examined *Ndrg1* and *Lcn2* expression levels in the pancreatic tissue of a genetically engineered mouse model of diet-induced PDAC (KRAS^G12D^/CRE) (5, 44). We showed that *Lcn2* expression was increased and *Ndrg1* expression was decreased in KRAS^G12D^/CRE mice, compared to the CRE control mice **(Figure 4D-4F).**

**Figure 4.** Iron treatment decreases expression of anti-metastatic marker NDRG1 in PDAC and *NDRG1* is inversely correlated with *LCN2* expression. (**A**) *NDRG1* expression levels in BXPC3 and PANC-1 under 150 µM FAC or 20 mM FAC treatments, relative to 0 mM FAC. One-way ANOVA test used to determine significance. (**B**) *NDRG1* expression levels in BXPC3 and PANC-1 under 20 µM DFO treatment, relative to 0 mM DFO. Unpaired t-test was used to determine significance. n=3-6 replicates (**C**) *LCN2,* and (**D**) *Lcn2* expression levels in a normal human pancreatic ductal epithelial cell line (HPDE), various human PDAC cell lines, and HPSC, relative to HPDE expression. (**E**) *NDRG1,* and (**F**) *Ndrg1* expression levels in mouse pancreatic tissue isolated from mice CRE and Kras^G12D^/CRE relative to expression in CRE control. n=3 replicates. One-way ANOVA test was used to determine significance in **B** and **D**; and unpaired t-test in **C** and **E**. Bars represent mean ± SEM. *p≤0.05, **p≤0.01, ***p≤0.001 ****p≤0.0001.

### Lcn2 depletion elevates *Ndrg1* expression, which is regulated in an iron-dependent manner

To further understand the role of LCN2 expression and iron levels in modulating NDRG1 expression in PDAC, we generated a KPC cell line with a biallelic *Lcn2* deletion. Several *Lcn2*^-/-^ (KO) clones were isolated and characterized by a series of genomic PCR assays, and two of those *Lcn2* KO clones were used in this study **(Figure 5 – figure supplement 1)**. Quantitative RT-PCR was performed to verify that no transcripts of *Lcn2* were present **(Figure 5 and Figure 5 – figure supplement 2A)**.

**Figure 5.**
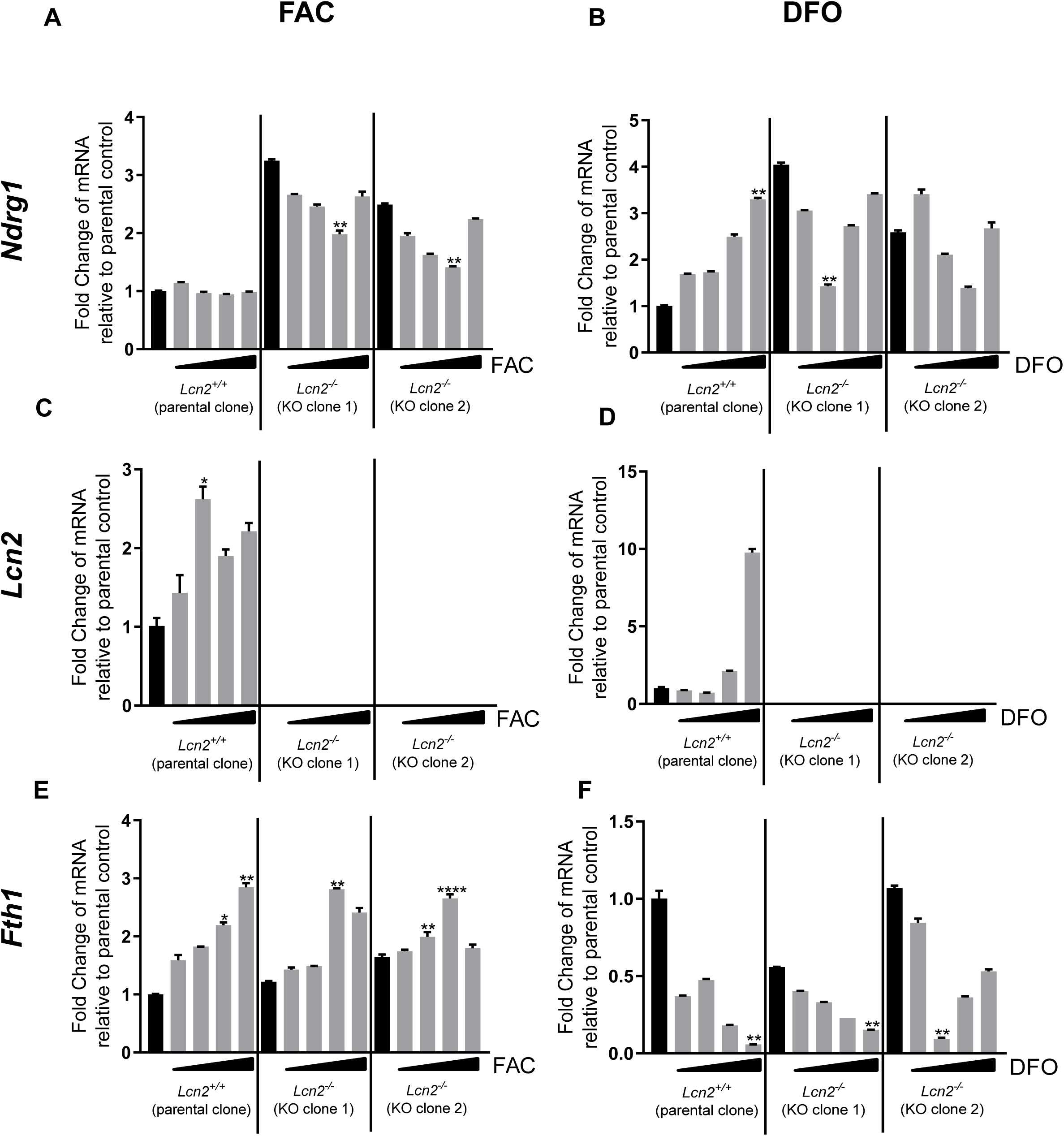
Lcn2 depletion elevates Ndrg1 expression, which is regulated in an iron-dependent manner. (**A**) *Ndrg1* expression in the *Lcn2*-KO clones after 24 hours of 0 µM, 25 µM, 50 µM, 150 µM, and 1500 µM FAC. (**B**) *Ndrg1* expression in mKPC controls and *Lcn2*-KO clones after DFO treatments of 0 µM, 1 µM, 10 µM, 50 µM, and 100 µM for 24 hours. (**C**) *Lcn2* expression in mKPC controls and *Lcn2*-KOs after same FAC treatments as Figure 5A. (**D**) *Lcn21* expression in mKPC controls and *Lcn2*-KOs after the same DFO treatments as Figure 5B. (**E**), *Fth1* expression in mKPC controls and *Lcn2*-KO clones after same FAC treatments as Figure 5A. (**F**) *Fth1* expression in mKPC controls and *Lcn2*-KO clones after the same DFO treatments as Figure 5B. n=3 replicates, Kruskal Wallis test was used to a determine significance. Bars represent mean ± SEM. *p≤0.05, **p≤0.01, ***p≤0.001 ****p≤0.0001.

Mouse PDAC cells (KPC-parental clone, *Lcn2*-KO clone 1 and *Lcn2*-KO-clone 2) were treated with similar FAC concentrations as the human PDAC cells shown in (**Figure 1)** and KPC cells prior to single cell cloning (**Figure 1 – figure supplement 2A**) to evaluate and compare cell proliferation. Results showed that FAC decreases proliferation in a dose-dependent manner for all KPC cell lines and at 72 hours, all KPC cell lines showed a decreased in proliferation similar to that observed for PANC-1 and HPSC in **Figure 1 (Figure 5 – figure supplement 2)**.

To investigate whether the lack of *Lcn2* expression modulates the levels of *Ndrg1* expression in cancer cells treated with iron, KPC-parental cells, *Lcn2*-KO clone 1 and *Lcn2*-KO clone 2 were treated with various concentrations of FAC and changes in *Ndrg1* gene expression were examined **(Figure 5A)**. We showed that *Ndrg1* expression was elevated 2.5 to 3.3-fold in the untreated *Lcn2*-KO clones relative to the parental clone KPC cell line. However, FAC treatments decreased the expression of *Ndrg1* in the *Lcn2*-KO clones and not the *Lcn2*^+/+^ KPC parental clone. Both *Lcn2*-KO clones showed a decrease in *Ndrg1* expression as the FAC concentration increased to 150 µM, but a reversal of this trend is observed at 1500 µM FAC for both KO clones **(Figure 5)**. The absence of *Lcn2* in mouse PDAC cells elevated the expression of *Ndrg1*, which was regulated by iron levels. To further understand this relationship, we measured *Lcn2* expression in the KPC-parental clone cells and found an overall increased in *Lcn2* expression at increasing doses of FAC **(Figure 5)**. These data suggest that iron levels regulate expression of *Lcn2* in cancer cells.

To verify iron influx into the cells, the expression of *Fth1* was quantified. *Fth1* expression was overall increased with increasing iron concentrations in all cells regardless of *Lcn2* expression **(Figure 5E)**. These results indicate that iron was being taken by the cells and stored even in the absence of *Lcn2* expression.

To assess whether iron chelation had the inverse effects of iron treatments on KPC cells with or without Lcn2 expression, we treated the cell lines with a range of DFO concentrations and measured the expression of *Ndrg1*. In the KPC-parental cells, increasing concentrations of DFO increased *Ndrg1* expression **(Figure 5B)**. However, the *Lcn2*-KO clones exhibited an overall decrease in *Ndrg1* expression. Furthermore, we measured *Lcn2* and *Fth1* expression after DFO treatments and showed that DFO increased *Lcn2* expression at high concentrations in the KPC-parental cell line **(Figure 5D)** and decreased *Fth1* expression overall in all KPC cell lines regardless of *Lcn2* expression **(Figure 5F)**. These results further depict the complexity of iron regulation and its association with other factors in PDAC. A plausible explanation for these results is that in a low iron environment, KPC cells decrease the expression of the iron storage gene *Fth1* because iron needs to be released from storage to be utilized by the cell, while increasing *Lcn2* expression in order to scavenge for iron and compensate for the lack of iron available in the cell.

### Lcn2 depletion regulates the expression of genes involved in extracellular matrix deposition and cell migration pathways

To further understand other pathways that are regulated by the lack of *Lcn2* expression in PDAC cells, we performed RNA sequencing analysis of the KPC parental and *Lcn2*-KO clone cell lines. A heat map of the hierarchal clustering of genes in KPC parental and *Lcn2*-KO clones shows that the clustering of genes differed greatly between the KPC parental and the *Lcn2*-KO clones, while the clustering of genes between both *Lcn2*-KO clones were similar to each other **(Figure 6A)**.

**Figure 6.**
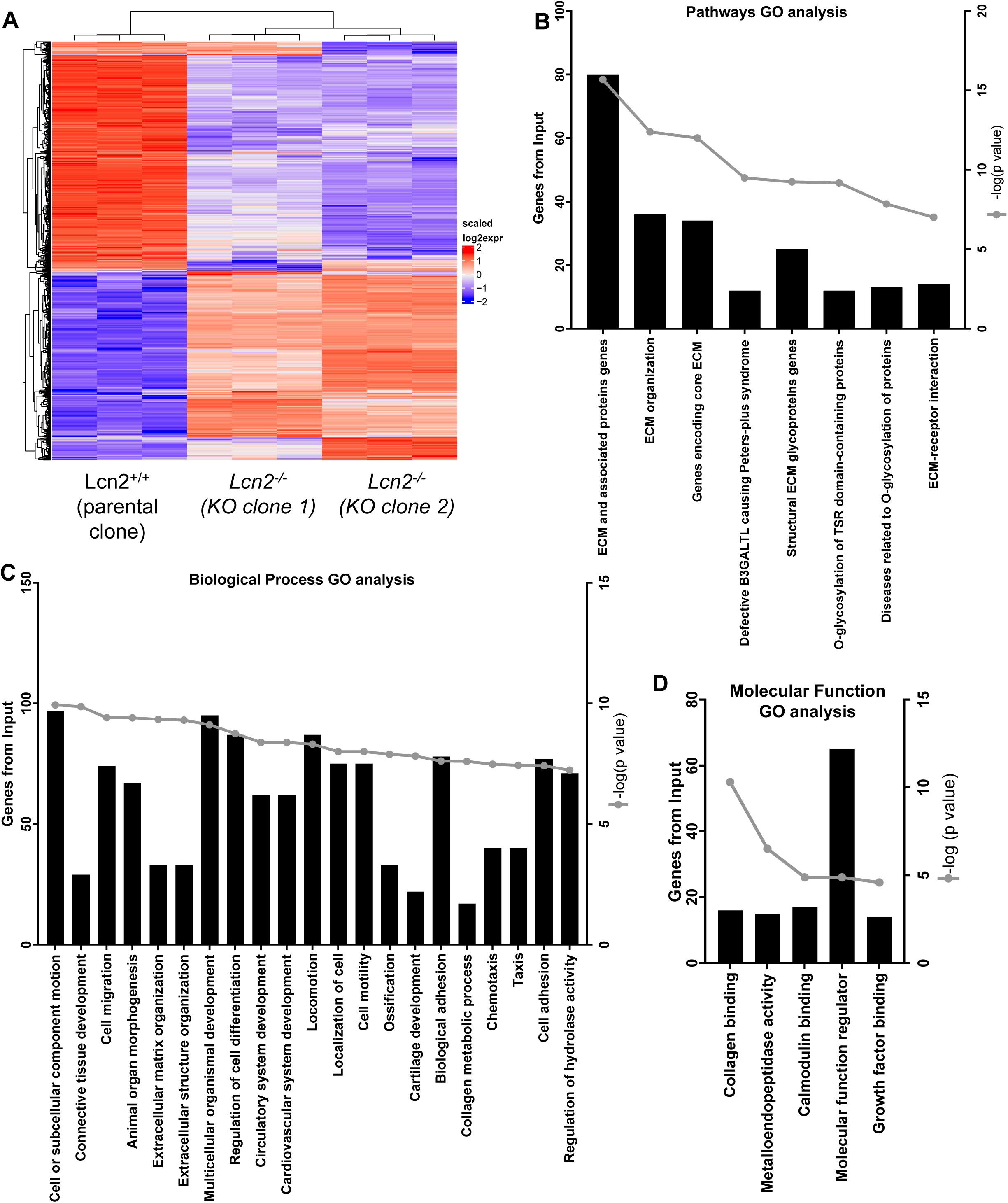
Lcn2 depletion regulates expression of Extracellular Matrix (ECM) related pathways. (**A**) Heat map. Hierarchical clustering of genes generated using R. The heatmap genes are colored proportional to voom log2 expression values. The color blue represents low expression of the respective gene, while the color red represents high expression. Changes from blue to red among the cell lines represent a relative increased in expression. Changes from red to blue among the cell lines represent a relative decrease in expression. (**B**) Gene ontology analyses of KPC parental and *Lcn2*-KO clone 1, showing the pathways associated with the genes differentially expressed in a *Lcn2*-KO clone. (**C**) Biological processes associated with genes differentially expressed in a *Lcn2*-KO clone. (**D**) Molecular function associated with the genes differentially expressed in a *Lcn2* KO.

Gene Ontology (GO) analyses were performed to identify the processes in which differentially expressed genes of these cell lines were involved. These results showed that *Lcn2* deletion differentially modulates ECM related mechanisms, cell migration, cell adhesion, blood vessel development, and connective tissue development, among others **(Figure 6B, C, D),** when compared to the KPC parental clone cell line expressing *Lcn2*. These are all characteristics involved in cancer development and progression, where LCN2 plays an important role (45, 46).

Furthermore, Gene Set Enrichment Analyses (GSEA) were used to identify classes of genes or proteins over-represented in the *Lcn2*-KO clones and to understand possible associations to cancer phenotypes. We found genes involved in the biological function of cell adhesion to be significantly over-represented in the KPC *Lcn2*-KO clone 1 **(Figure 7 A)** and clone 2 **(Figure 7 B)** cell lines, compared to the KPC parental clone. Genes involved in proteinaceous extracellular matrix processes and the integrin pathway were also found significantly over-represented in the KPC *Lcn2*-KO clone 1 **(Figure 7 C, E)** and clone 2 **(Figure 7 D, F)** cell line compared to KPC parental. Therefore, *Lcn2* deletion modulates a large number of genes involved in extracellular matrix deposition and cell migration pathways.

**Figure 7.**
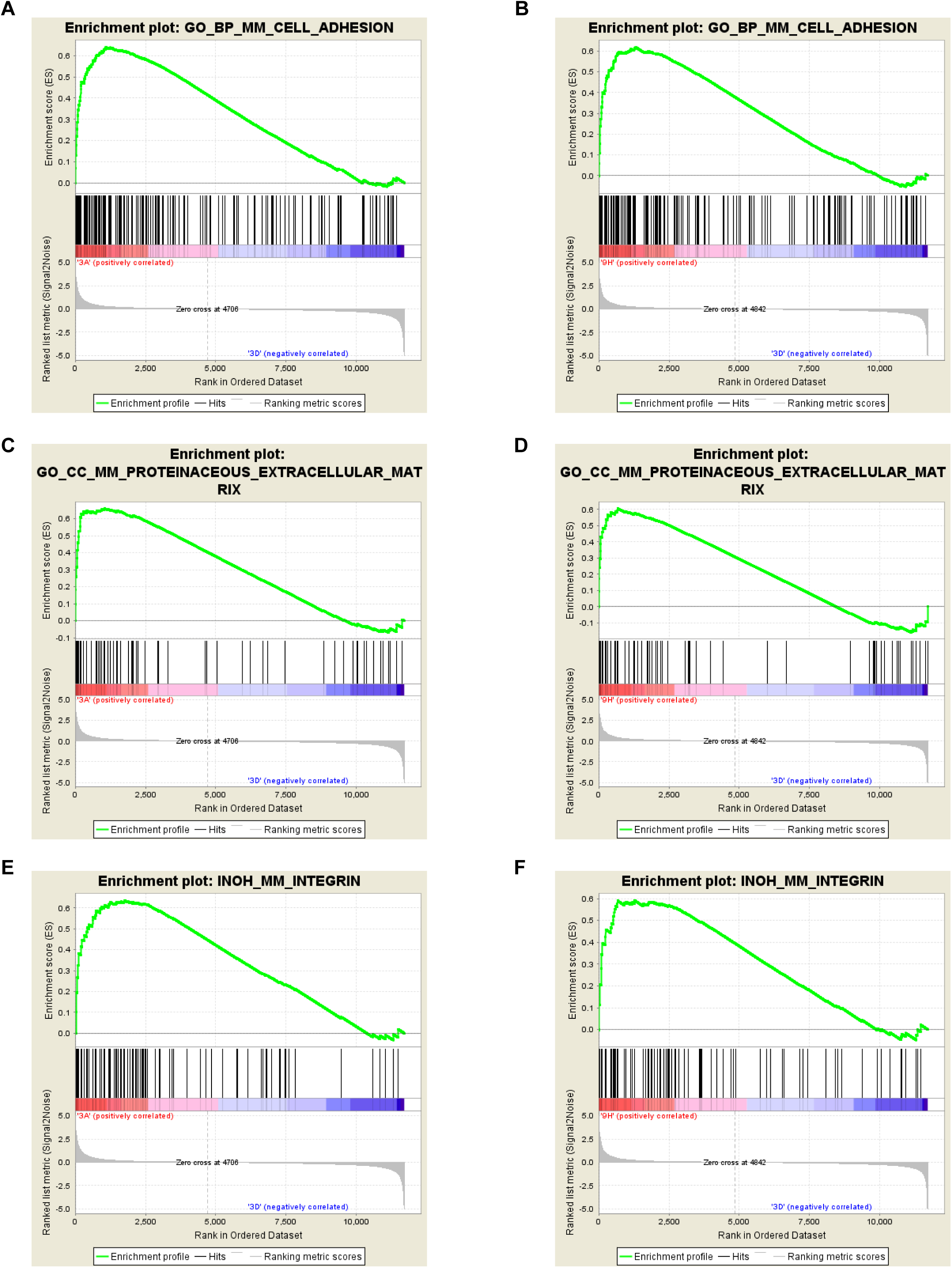
Lcn2 depletion regulates expression of Extracellular Matrix (ECM) related pathways. GSEA analysis showing overrepresentation of (**A**) cell adhesion; (**C**) proteinaceous extracellular matrix, and (**E**) integrin pathway in mKPC *Lcn2*-KO clone 1 vs mKPC parental clone. (**B**) cell adhesion; (**D**) proteinaceous extracellular matrix, and (**F**) integrin pathway in mKPC *Lcn2*-KO clone 2 vs mKPC parental clone.

## DISCUSSION

Since iron is essential for cell growth, DNA synthesis, and apoptosis, tumor cells tend to have elevated iron requirements relative to normal somatic cells due to their increased proliferation rates (47). In this study, the human PDAC cell line BXPC3, increased proliferation in response to changes in iron concentration compared to PANC-1 cells **(Figure 1)**. BXPC3 cells express an un-mutated *KRAS* gene while PANC-1 express a mutant *KRAS^G12D^* (37, 48, 49), possibly accounting for the differences observed in these cell lines. These effects are slightly reversed when iron is chelated with DFO. Interestingly, in PDAC patients in which Kras is not mutated, iron concentrations are significantly higher (50). This suggests that Kras mutation status could be involved in iron homeostasis in the same way that the TP53 mutation, another common mutation in PDAC patients, was found to regulate cellular iron transport and storage (51). Moreover, triapine, an iron chelator and an inhibitor of the M2 subunit of the ribonucleotide reductase, has been used to improve radiation therapy outcomes in PDAC patients (52). In addition, iron chelation has also been effective in reducing tumor growth alone or in combination with other treatments in vitro and in PDAC xenograft models (53–56).

Considering the interaction of iron with the pro-inflammatory cytokine LCN2, we sought to understand whether iron contributed to the inflammatory responses observed in PDAC. Here we showed that iron induced inflammation by increasing the expression of *IL6* and *IL1β* in both PDAC cancer and pancreatic stellate cells. However, the expression of the LCN2 receptor *SLC22A17* remained unchanged **(Figure 2 A, C, G)**. In breast cancer, iron contributes to chemoresistance by increasing IL6 production in tumor associated macrophages (57). Chemoresistance is one of the most common features in PDAC in part due to the dense stromal environment (58). For this reason, additional studies should investigate whether iron modulation could be used in combination with current therapies to decrease chemoresistance in PDAC.

Iron concentrations are associated with EMT in tumors cells (10, 11). The crucial steps of EMT leading to metastasis are characterized by a decrease in intercellular adhesion of the tumor cells, a loss of epithelial morphology, and increased invasion, a hallmark of mesenchymal morphology (27, 59). At the molecular and transcriptional level, EMT is characterized by promoting the degradation of basement membranes and ECM, leading to invasion and metastasis (46). Here we showed that excess iron in the media of BXPC3 cells increased the expression of EMT-associated markers and cell invasion more than PANC-1 cells **(Figure 3)**, while iron chelation decreased mesenchymal markers expression in both cell lines (**Figure 3 - figure supplement 1A, B**). Differences in the Kras mutation status between BXPC3 and PANC-1 cell lines could be mediating the invasive phenotype and proliferation that results from iron in PDAC (35, 37, 48). Previous reports suggest that *Ras* expression can modulate cellular processes such as cell survival in ovarian cancer (36) and other factors in PDAC (50, 60). Iron chelation has been effective at suppressing EMT in lung, prostate, colon and esophageal cancer (20–23). However, a study in mice showed that while EMT induces chemoresistance in PDAC, EMT is not needed to develop an invasive and metastatic phenotype (61). Contributing to the Kras expression difference among the cell lines, we found higher *LCN2* expression in BXPC3 cells compared to the mutant *KRAS* PANC-1 cells **(Figure 4C)**, which might support the differences in iron responses and iron-dependent EMT initiation between the lines. Future studies aimed at understanding whether LCN2 expression is associated with Kras mutations in PDAC are necessary to determine whether LCN2 and iron targeted therapy will benefit PDAC patients.

Besides inducing EMT, iron also regulates expression of the metastatic suppressor NDRG1 (21, 23, 34). NDRG1 expression can be regulated by other effectors including N-myc, acetylation of histones, hypoxia, and intracellular calcium levels (21, 34, 41, 42, 62, 63). NDRG1 is responsible for the suppression of glycolytic metabolism in PDAC, a metabolic pathway utilized by many cancers for growth (60). In our study, we found that increased levels of iron downregulated the expression of *NDRG1*, while iron chelation upregulated *NDRG1* expression in PDAC cancer cells. Moreover, baseline expression of *NDRG1* was lower in the PANC-1 cell line compared to the BXPC3 cell line **(Figure 4A, B).** Similar results were observed in the mouse pancreatic cancer cell line, where the induction of oncogenic Kras^G12D^ mutation decreased significantly the protein levels of NDRG1 (60). Since *NDRG1* expression is already reduced in the mutant Kras compared to wild type Kras cell line, the further decrease in *NDRG1* expression caused by iron might not have the same impact on promoting EMT. Although Kras mutations and elevated expression of LCN2 are common in PDAC, our data suggests that knowing the mutation status of Kras and LCN2 levels in patients could help inform whether iron modulation and LCN2 blockade could serve as a novel treatment approach.

Intracellular iron is regulated by the expression of FTH1, which functions as an iron storage protein and controls intracellular iron release (64). Here we showed that FTH1 expression is lower in PDAC cell lines compared to a normal human pancreatic ductal epithelial cell line **(Figure 3 - figure supplement 1)**. These data suggest that there is an increase of free iron and possible reduction in iron storage due to the high demand for iron by PDAC cells. *FTH1* expression is also downregulated in HPSC, potentially for the same reason. Lower amounts of intracellular FTH1 result in higher concentrations of free iron, contributing to elevated reactive oxygen species production, another essential factor in EMT (26). In human breast and lung cancer cell lines, inhibition of *FTH1* expression promoted migration, decreased adhesion, and displayed a fibroblastoid morphology that lead to EMT (19). These findings support the observation that increased extracellular and intracellular iron, as well as reduced intracellular FTH1, results in EMT.

To further understand how LCN2 expression in cancer cells modulated iron responses, LCN2 expression was deleted via CRISPR. RNA sequencing was used to identify other processes regulated by *Lcn2* expression in PDAC. We showed that *Lcn2* deletion is associated with differentially expressed genes involved in extracellular matrix related pathways and processes vital to cancer progression, such as cell migration, cell motility, and collagen binding (**Figure 6 B, C, D**). These results validate and build upon our prior work supporting the finding that stromal cells treated with LCN2 increase inflammation in the TME of PDAC and that lack of *Lcn2* expression in mice delays PDAC tumor formation and increases survival (5). In particular, the processes of cell adhesion, proteinaceous extracellular matrix, and integrin pathway were strongly over-represented in the tumor cells that lack *Lcn2* expression **(Figure 7**). Cell adhesion is associated with an epithelial phenotype and it is known to be reduced during EMT and metastasis (27). Moreover, the proteinaceous extracellular matrix and integrin pathways are tightly associated with the organization of the extracellular matrix in cancer progression (65).

Overall, in our study we demonstrated that iron promotes inflammation, invasion, and EMT on PDAC cell lines partially by modulating NDRG1 expression. Iron responses were strongly associated with LCN2 expression, which contributes to the formation and function of the TME. Finally, our data suggests that iron chelation can potentially decrease invasion and metastasis in PDAC, especially if combined with an LCN2 blockade. Additional studies are needed to assess the therapeutic potential of LCN2 blockade and iron modulation on PDAC progression and metastasis, and its correlation with *Kras* mutations status. Our study also suggests that *Kras* mutations may play a role in how cancer cells respond to iron in PDAC. Therefore, knowing the mutation status of *KRAS* in PDAC would help determine whether a patient could benefit from iron chelation therapy and LCN2 blockade in conjunction with current treatment standards of care.

## Materials and Methods

### Cell culture and cell lines

Cell lines were cultured at 37°C with 5% CO_2_ in DMEM supplemented with 4.5 g/l glucose, L-glutamine and 10% FBS. All cell lines tested negative for the presence of mycoplasma using MycoAlert kit (Lonza, Hayward, CA), which we test monthly. BXPC3, PANC-1, HPAC, CAPAN2, MIAPACA2, CAPAN1, and MPANC96 were obtained from the American Type Culture Collection (ATCC). HPDE (non-malignant, human pancreatic ductal epithelial) cells (66, 67) were obtained from Dr. Tsao (Ontario Cancer Institute, Toronto, ON, Canada). Mouse PDAC cells were derived from a pancreatic tumor of a *LSL-KRas^G12D^*, *LSL-Trp53^-/-^*, *PDX-1-CRE*, (KPC) genetically engineered mouse, as described in (5, 68). The KPC cell line was subcloned to derive a single clone for CRISPR cloning. HPSC were acquired from a resected human PDAC sample as described in (5).

### CRISPR plasmids, guide RNA constructs and oligonucleotides

The PX459V2.0 plasmid (pSpCas9(BB)-2A-Puro) (Addgene, Watertown, MA) was used to ligate the appropriate single guide RNA sequences 5’ of the trans-activating CRISPR RNA (tracrRNA) scaffold to create a gRNA/tracrRNA that will direct the Cas9 nuclease to the appropriate site for cleavage (69). Cloning of single guide RNA sequences, transfection of plasmids into cells, and selection of biallelic, *Lcn2*-deleted clones are modified from previously described protocols (70). PX459V2.0 plasmid contains the *cas9* ORF, the tracrRNA scaffold ORF, a cloning site (BbsI) for ligation of a specific guide RNA at the 5’ end of the tracrRNA scaffold ORF, and a puromycin resistance gene for selection of cells that were transfected with PX459V2.0. Transfection of PX459V2.0 was performed using Lipofectamine 3000 (Invitrogen) under manufacturer’s guidelines. Guide RNA sequences are located in intron 2 (gRNA-1), intron 5 (gRNA-2), and the 3’ UTR (gRNA-3) of the mouse *Lcn2* gene (six exons total) (**Figure 5 – figure supplement 1**). Paired guide RNAs, gRNA-1/gRNA-2 and gRNA-1/gRNA-3, generated deletions from exons 3 through 5 and exons 3 through 6, respectively, thus creating two distinguishable types of genomic deletions. Biallelic *Lcn2*-deleted single cell-derived clones were identified using genomic PCR and oligonucleotides 5’ to gRNA-1 genomic location, and 3’ to gRNA-2 and gRNA-3 genomic locations to detect deletions in the *Lcn2* locus (**Figure 5 – figure supplement 1**). Biallelic *Lcn2*-deleted clones were further verified using oligonucleotides internal to the deleted region (**Figure 5 – figure supplement 1**).

Guide RNA oligonucleotide pairs for ligation into PX459V2.0 and Genomic PCR oligonucleotides are displayed in **Supplementary file 1**.

### Iron treatments

For iron and DFO treatments, plated cells were washed with PBS three times and serum starved 24 hours prior to the treatments. Treatments with iron were performed with Ferric Ammonium Citrate (FAC) (Sigma-Aldrich, St. Louis, MO), in concentrations ranging from 0-20mM FAC added to the serum-free media (SFM) (only DMEM) for 24-72 hours depending on the experiment. Iron chelation was performed with DFO (Sigma-Aldrich, St. Louis, MO) in concentrations ranging from 0-50µM added to SFM for 24-72 hours depending on the experiment.

### Cell proliferation

CellTiter 96 AQueous One Solution Cell Proliferation Assay MTS 3-(4,5-dimethylthiazol-2-yl)-5-(3-carboxymethoxyphenyl)-2-(4-sulfophenyl)-2H-tetrazolium) was used according to the manufacturer’s protocol (Promega, Madison, WI) to assess proliferation of cells cultured in 96-well plates.

### Cell invasion

Invasion assays were performed according to the manufacturer’s protocol with some modifications. Briefly, the upper surface of transwell membranes (24-well plates, 6.5mm Insert, 8.0um PET membrane; (Trevigen) was coated with Basemen Membrane Extract (BME), composed of laminin I, collagen IV, entactin and heparin sulfate proteoglycan. The coated transwells were incubated at 37^0^ C, 24 hours before the assay. The following day, SFM or 10% FBS DMEM was added to the bottom chambers. The cells were centrifuged and resuspended in 0.25 mg BSA/PBS twice, and 50,000 cells were added to the upper transwell membrane chambers. The cells were left to invade overnight into either SFM or 10% FBS DMEM, at 37^0^ C. The next day, the cells that migrated to the bottom were washed and stained with Calcein-AM/cell dissociation solution and incubated for 1 hour. Cell numbers were read at 485 nm excitation and 520 nm emission wavelengths using the Synergy HT multimode micro-plate reader (BioTek, Winooski, VT).

### DNA isolation and genomic PCR

DNA from KPC cultured cells, for the selection of CRISPR-derived biallelic *Lcn2* deletion clones, was isolated using the DNeasy Blood & Tissue kit (Qiagen, Venlo, Netherlands). Genomic PCR was performed using 30 ng of genomic DNA for the detection and verification of the biallelic deletion within the mouse *Lcn2* gene.

### RNA isolation and quantitative RT-PCR

RNA isolation from cultured cells or mouse pancreatic tissue was performed using TRIzol reagent (Life Technologies, Carlsbad, CA) following the manufacturer’s protocol. Reverse transcription to generate cDNA from total RNA was performed using the Verso cDNA synthesis kit (ThermoFisher Scientific, Waltham, MA). Quantitative PCR using TaqMan primers (**Supplementary file 1**) (ThermoFisher Scientific, Waltham, MA) was employed to determine gene expression levels compared to control and normalized to either 18S or GAPDH.

### Imaging

Bright-field microscopy images of BXPC3, PANC-1 and HPSC cell lines were taken with an Olympus IX51 microscope DP74 digital camera.

### Genetically engineered transgenic mice and treatments

*KRas*^G12D^ mice obtained from the Mouse Models of Human Cancer Consortium Repository (NIH Bethesda, MD) (71) were bred with the Ela-CreERT (CRE) mice as previously described (72) to generate the *KRas*^G12D^/CRE mice. At 40 days old, mice were administered tamoxifen orally for 3 consecutive days and were fed a high fat diet for 6 weeks (Test Diet DIO 58Y1 van Heek Series; Lab Supply, Fort Worth, TX), in which 60% of energy was derived from fat. Pancreatic tissue was collected after the intervention and RNA was extracted.

### RNA profiling

The RNA expression was analyzed by RNAseq. The RNA library was generated using the NEB Next Ultra II Directional RNA kit. The sequencing approach was polyA-selection (mRNA-seq). The input amount was 200 ng total RNA as determined by ThermoFisher/LifeTechnology Qubit RNA assay. Each library was sequenced to a depth of 17 – 20 million passed filter clusters (or 34 – 40 million passed filter reads) using the Illumina HiSeq 4000 sequencer paired – end 150bp approach. The data discussed in this publication have been deposited in NCBI’s Gene Expression Omnibus (73) and are accessible through GEO Series accession number GSE143463 (https://www.ncbi.nlm.nih.gov/geo/query/acc.cgi?acc=GSE143463).

### RNAseq data processing and analysis

Sequencing reads were aligned to mouse reference genome GRCm38 with hisat2 (74). Gene expression was quantified with featureCounts software (75) for genes annotated by ensembl Mus_musculus.GRCm38.83, counting the primary alignment in the case of multimapped reads. Genes were included if at least half of the samples had an expression of 2 CPM. Raw counts were normalized by voom and differential expression was performed with limma (76).

For the heatmap, we selected genes with logFC > 1 or <-1 and with FDR<0.05 in either KPC parental clone vs KPC *Lcn2*-KO clone 1 or KPC parental clone vs KPC *Lcn2*-KO clone 2. Using ComplexHeatmap in R, we plotted scaled voom normalized expression values (77). To identify gene sets enriched in pairwise comparisons of sample groups, we performed Gene Set Enrichment Analysis (GSEA) (78, 79). For the analysis, we used voom-normalized expression data. Mouse gene sets were downloaded from Gene Set Knowledgebase GSKB (80) for Gene ontology (GO), pathway, and transcription factors.

### Statistics

Statistical analyses were carried out using the Prism 5 software program (GraphPad Software San Diego, CA). Data are presented as the mean ± standard error of the mean. A t-test or one-way analysis of variance (ANOVA) was performed to analyze the data for two groups and multiple comparisons respectively. Multiple comparisons were corrected with the Dunnett’s test. Levels of significance are indicated as follows: ∗ = *P* ≤ 0.05, ∗∗ = *P* ≤ 0.01, ∗∗∗ = *P* ≤ 0.001, ∗∗∗∗ = *P* ≤ 0.0001. Non-parametric analyses were conducted when data were not normally distributed. A minimum of three replicates were performed for all in-vitro studies.

## AUTHOR CONTRIBUTIONS

1. Valentina Pita MS - study concept and design; development of methodology; acquisition of data; analysis and interpretation of data; drafting of initial manuscript; writing, review, and/or revision of the manuscript; administrative, technical, or material support; final approval of the version to be submitted;
2. Andrew W. Dangel PhD - study concept and design; development of methodology; acquisition of data; analysis and interpretation of data; drafting of initial manuscript; writing, review, and/or revision of the manuscript; administrative, technical, or material support; final approval of the version to be submitted;
3. Kristyn Gumpper PhD – writing, review, and revision of the manuscript, final approval of the version to be submitted;
4. Andrea Ludwig - development of methodology; acquisition of data, analysis and interpretation of data, administrative, technical, or material support, final approval of the version to be submitted;
5. Olivia Ueltschi - acquisition of data, analysis and interpretation of data, final approval of the version to be submitted;
6. Xiaokui Mo PhD - acquisition of data, analysis and interpretation of data, final approval of the version to be submitted;
7. Maciej Pietrzak PhD - acquisition of data, analysis and interpretation of data, final approval of the version to be submitted;
8. Amy Webb PhD - acquisition of data, analysis and interpretation of data, final approval of the version to be submitted;
9. Rosa F. Hwang MD - administrative, technical, or material support; final approval of the version to be submitted;
10. Madelyn Traczek - development of methodology; acquisition of data, analysis and interpretation of data, administrative, technical, or material support, final approval of the version to be submitted;
11. Niharika Badi, MS - development of methodology; acquisition of data, analysis and interpretation of data, administrative, technical, or material support, final approval of the version to be submitted;
12. Zobeida Cruz-Monserrate, PhD - study concept and design; development of methodology; acquisition of data; analysis and interpretation of data; drafting of initial manuscript; writing, review, and/or revision of the manuscript; administrative, technical, or material support; final approval of the version to be submitted; study supervision.

## Acknowledgements

We thank The Ohio State University Genomics Shared Resource (GRS) for the RNA sequencing library generation which is funded by NCI Cancer Center Support Grant P30 CA016058.

---

**Figure 1 – figure supplement 1.**
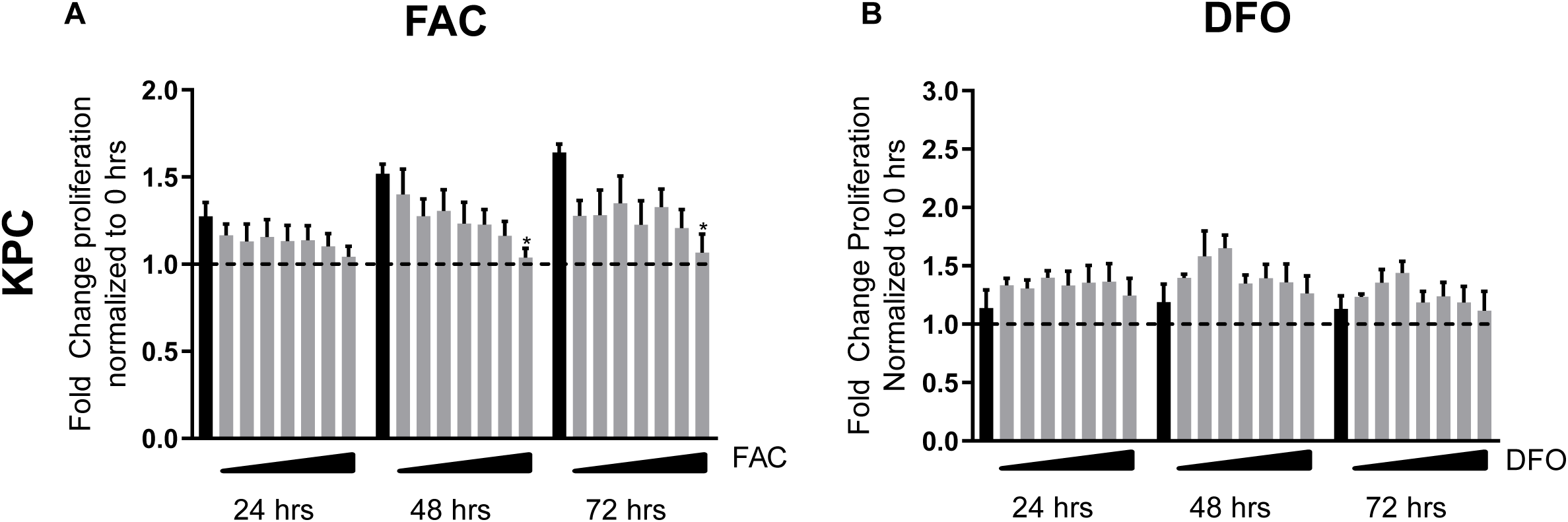
Iron and DFO treatments regulate cell proliferation and viability in the TME. (**A**) Mouse PDAC cell line KPC treated with same concentration of FAC as Figure 1A. (**B**) Mouse PDAC cell line KPC treated with same concentration of DFO as Figure 1B. Significance was assessed by one-way ANOVA. Bars represent mean ± SEM. *p≤0.05, **p≤0.01, ***p≤0.001 ****p≤0.0001. n=3-6 per group.

**Figure 3 - figure supplement 1.** Gene expression levels for *ZEB1*, *SNAI1*, *TWIST*, and *CDH1* in BXPC3 (**A**) and PANC-1 (**B**) relative to 0 mM DFO (-) treatment, (+) denotes 20 µM DFO. Treatments were maintained for 48 hours. n≥3. (**C**) *FTH1* expression levels in the same cell lines as denoted in (Figure 4.A). Significance was assessed by one-way ANOVA. Bars represent mean ± SEM. *p≤0.05, **p≤0.01, ***p≤0.001, ****p≤0.0001. n=3 replicates per group.

**Figure 5 – figure supplement 1.**
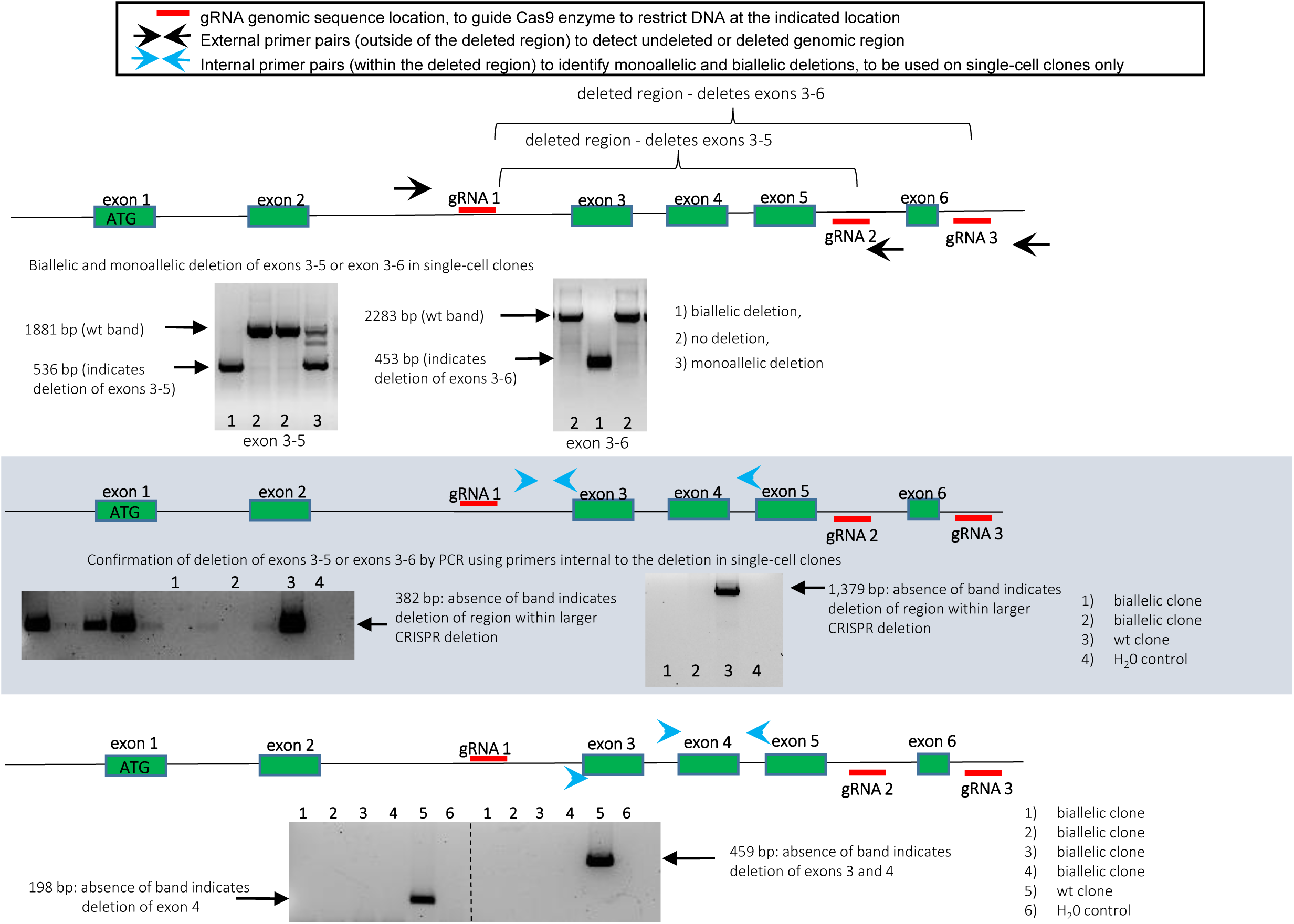
Mouse *Lcn2* gene – gRNA placement and PCR detection of deleted regions via CRISPR.

**Figure 5 – figure supplement 2.**
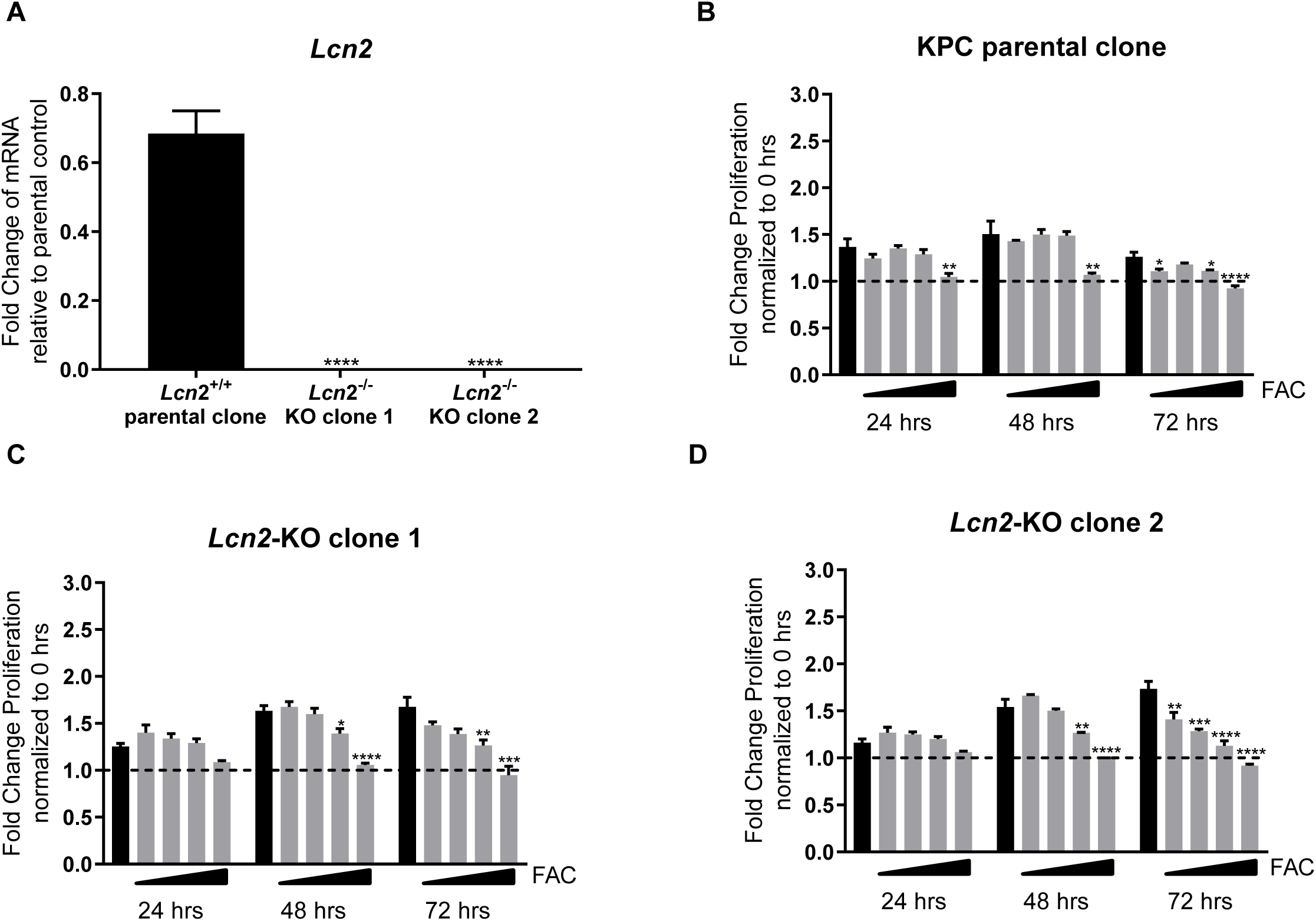
Two distinct clones of a CRISPR-derived biallelic *Lcn2* deletion in KPC. (**A**) Gene expression levels for *Lcn2* in KPC parental clone, *Lcn2*-KO clone 1 and KPC *Lcn2-*KO clone 2 relative to KPC parental clone. (**B**) Iron treatments affect cell proliferation and viability in KPC parental clone. (**C**) *Lcn2*-KO clone 1, and (**D**) *Lcn2*-KO clone 2. Cells were treated with 0 mM, 1.5 mM, 5 mM, 10 mM, and 20 mM FAC over 72 hours. Results were normalized to 0 hours, represented by a horizontal dashed line. Significance was assessed by one-way ANOVA. Bars represent mean ± SEM. *p≤0.05, **p≤0.01, ***p≤0.001 ****p≤0.0001. n=3 replicates per group.

**Supplementary File 1.** Guide RNA pairs for ligation into PX459V2.0 and Genomic PCR oligonucleotides for CRISPR-derived biallelic *Lcn2* deletion in KPC and list of TaqMan primers used in the qPCR expression measurements.

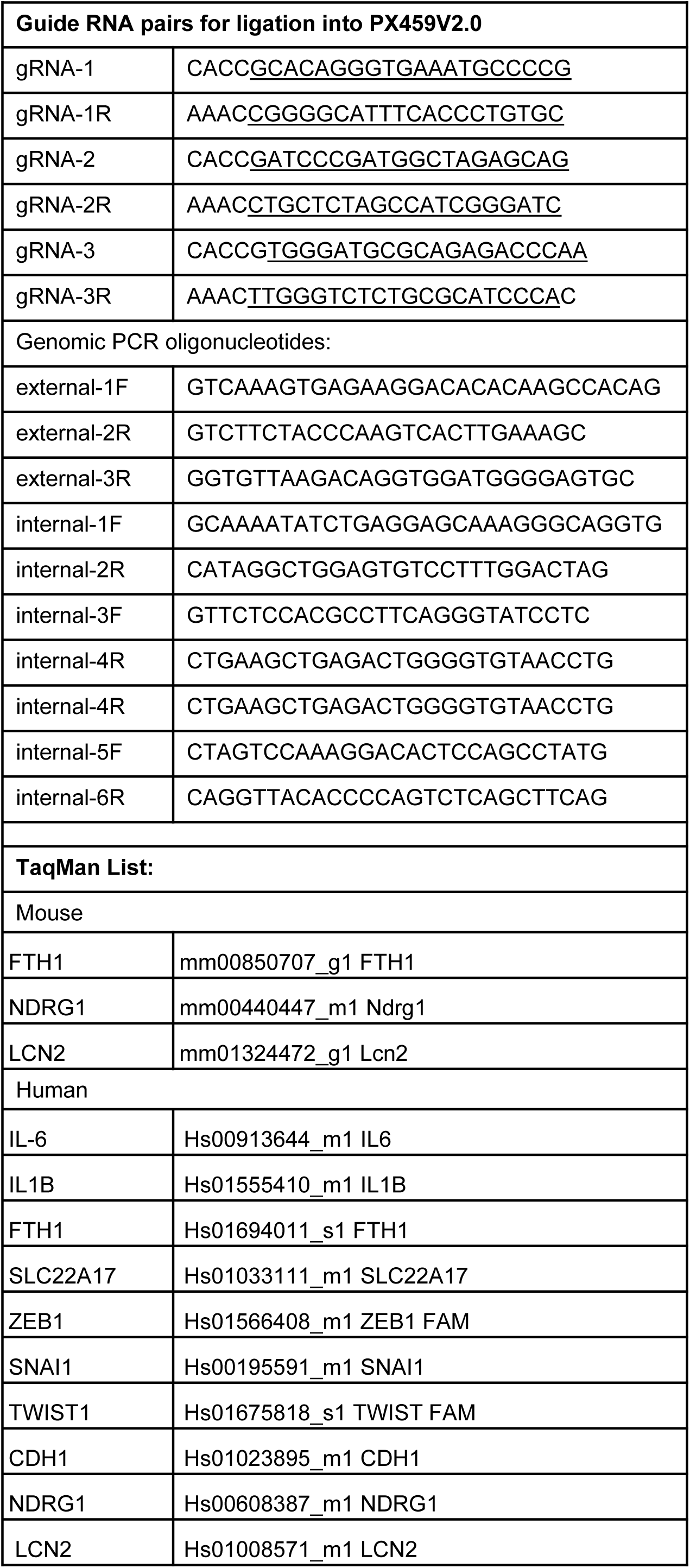

